# The GSA Family in 2025: A Broadened Sharing Platform for Multi-Omics and Multimodal Data

**DOI:** 10.1101/2025.03.31.646270

**Authors:** Sisi Zhang, Xu Chen, Enhui Jin, Anke Wang, Tingting Chen, Xiaolong Zhang, Junwei Zhu, Lili Dong, Yanling Sun, Caixia Yu, Yubo Zhou, Zhuojing Fan, Huanxin Chen, Shuang Zhai, Yubin Sun, Qiancheng Chen, Jingfa Xiao, Shuhui Song, Zhang Zhang, Yiming Bao, Yanqing Wang, Wenming Zhao

## Abstract

The Genome Sequence Archive family (GSA family) provides a suite of database resources for archiving, retrieving, and sharing multi-omics data for the global academic and industrial communities. It currently comprises four distinct database members: the Genome Sequence Archive (GSA, https://ngdc.cncb.ac.cn/gsa/), the Genome Sequence Archive for Human (GSA-Human, https://ngdc.cncb.ac.cn/gsa-human/), the Open Archive for Miscellaneous Data (OMIX, https://ngdc.cncb.ac.cn/omix/), and the Open Biomedical Imaging Archive (OBIA, https://ngdc.cncb.ac.cn/obia). Compared to the 2021 version, the GSA family has significantly expanded by introducing a new database member, the OBIA, and through comprehensive upgrades to the existing ones. Notable enhancements to the existing members include broadening the range of accepted data types, enhancing the quality control systems, fine-tuning the data retrieval system, and refining the data-sharing management mechanisms.

## Introduction

Next-generation sequencing (NGS) technologies have revolutionized genomics, leading to an exponential increase in the amount of available raw sequence data and the expansion of researchers’ understanding of genome structure, function, and complexity [1, 2]. The advancements in high-throughput omics technologies have empowered researchers to adopt multi-omics integration strategies to integrate and analyze data from different layers, including genomics, transcriptomics, epigenomics, proteomics, and metabolomics, thereby deepening our understanding of complex biological phenomena [3-5]. In recent years, advances in medical imaging technologies—such as Magnetic Resonance Imaging (MRI), Computed Tomography (CT), and Positron Emission Tomography (PET)—along with improvements in image analysis, have enabled the extraction of detailed imaging features. These features can now be integrated with multi-omics data, opening new avenues for medical research and personalized healthcare [6-8]. These factors collectively have driven the transition from single-omics to multi-omics research and have advanced data management toward standardization, normalization, and platform-based sharing. The management and sharing of multi-omics and multimodal data present unique complexities, primarily in the following aspects: the large scale of data results in significant storage and processing challenges, requiring efficient solutions; the diverse sources and varying formats of data necessitate the implementation of flexible management strategies; and the imperative for data security and privacy protection demands the enforcement of stringent management measures. Therefore, a robust data management platform is needed to deliver comprehensive solutions for efficient storage, rapid retrieval, and secure sharing of the multi-omics and multimodal data in life sciences, to advance the progress of scientific research and clinical applications.

The Genome Sequence Archive [9] serves as a repository for raw sequence data in the National Genomics Data Center [10] at China National Center for Bioinformation (CNCB-NGDC)[10, 11], which has effectively addressed the longstanding challenges related to the unified submission and management of genomic data in China, promoting the sharing and reuse of data within the community. In 2021, the GSA family [12], comprising GSA, Genome Sequence Archive for Human (GSA-Human), and the Open Archive for Miscellaneous Data (OMIX), was introduced, greatly enhancing the management and sharing capabilities for multi-omics data.

Over the past few years, the GSA family has made significant efforts to address the challenges posed by the rapid accumulation of multi-omics and multimodal data. Firstly, the GSA family has expanded the range of data types it handles, adding biomedical imaging and clinical data, which are now archived in a new member database called the Open Biomedical Image Archive (OBIA) [13]. Through this platform, researchers can obtain and integrate data from various domains more comprehensively. In particular, by combining raw sequence data with clinical and medical imaging data, researchers are able to delve deeper into the molecular mechanisms of diseases, providing crucial support for the development of personalized treatment plans. Secondly, the GSA family has carried out a comprehensive upgrade of the core functions of its existing databases, including enhancements to data quality control, search tools, data sharing mechanisms, and security measures. These improvements not only increase the accuracy and efficiency of data processing but also ensure the convenience and security of data sharing and access. In particular, the optimization of the quality control process has significantly improved the precision and reliability of the data, while also standardizing the archived data, thus providing higher-quality corpus resources for AI-driven data analysis. Additionally, the GSA family has mirrored and integrated raw sequence data from the National Center for Biotechnology Information (NCBI), providing unified retrieval and download services for the submitted and integrated data. This initiative has made cross-platform data integration smoother, facilitated the sharing and analysis of large-scale datasets, and supported research and innovation in the life sciences.

### Data model

In 2024, the GSA family has expanded to four member databases: GSA, GSA-Human, OMIX, and OBIA. These databases are tightly integrated through BioProject and BioSample (**Figure 1**), forming a comprehensive multi-omics data framework that ensures seamless data sharing.

**Figure 1.**
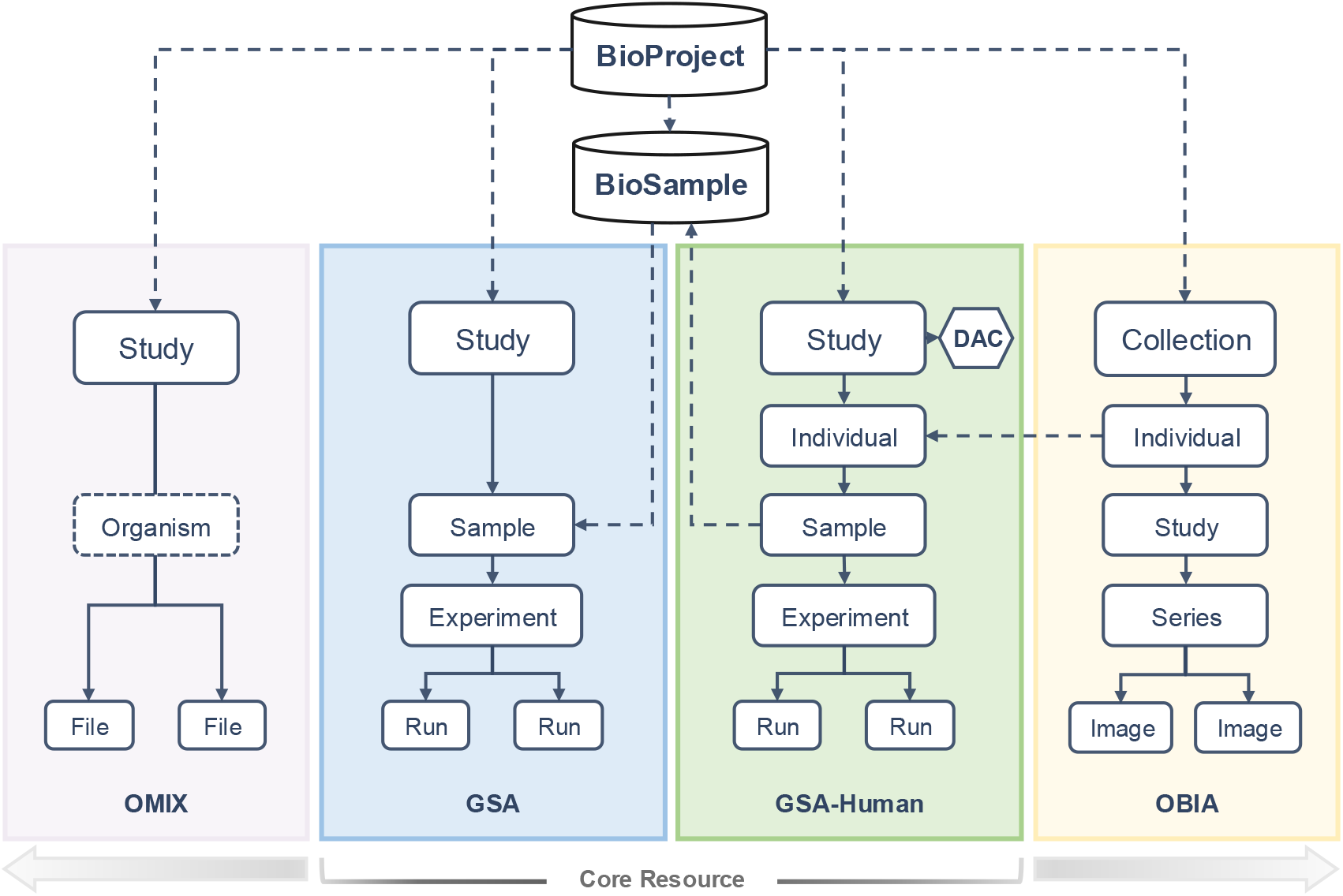
GSA family data model. BioProject and BioSample are two independent meta-information databases, which offers structured storage for research purposes and biomaterials, respectively. All members of the GSA family are linked to BioProject, thereby creating a comprehensive framework for multi-omics data. GSA-Human established a data synchronization process with BioSample. Individuals in OBIA can be linked to GSA-Human by accessions. OMIX is for various types of data that are unsuitable for submission to other data family members. DAC, Data Access Committee.

In GSA, metadata is organized into four objects: Study, Sample, Experiment, and Run. Each study is identified by a BioProject (https://ngdc.cncb.ac.cn/bioproject), a key meta-information database that provides structured storage for program descriptions. And each sample is assigned a unique BioSample accession number, with metadata systematically archived and retrievable through the BioSample database (https://ngdc.cncb.ac.cn/biosample).

GSA-Human employs a structured framework comprising five core objects: Study, Individual, Sample, Experiment, and Run. Each research study is uniquely identified by a BioProject accession. While GSA-Human does not directly utilize the BioSample object for sample representation, it maintains synchronization of non-sensitive sample metadata with the BioSample database. This integration facilitates the seamless sharing of sample metadata both within the GSA family and across CNCB-NGDC databases, such as Genome Warehouse[16] and Genome Variation Map[17], through BioSample accessions.

OBIA includes five key objects: Collection, Individual, Study, Series, and Image. Notably, Individuals within this framework can be linked to GSA-Human via individual accessions, streamlining the integration of imaging and raw sequencing data for multi-omics research.

OMIX uses a lightweight data model with three objects: Study, Organism, and File, providing flexibility and scalability to accommodate diverse researcher submission need. By integrating with external datasets through BioProject accessions, OMIX serves as a key complement of the GSA family. It supports specialized scientific data types that are related to but beyond the reception scope of other member databases.

### Archival resources

#### Genome Sequence Archive

GSA, built based on the International Nucleotide Sequence Database Collaboration (INSDC) [14] data standards, is an open-access repository dedicated to archiving, retrieving, and sharing non-human raw sequence data. In 2023, GSA was selected as one of the Global Core Biodata Resources by the Global Biodata Coalition, demonstrating its pivotal role in the global landscape of biological data resources.

To enhance the user experience and streamlined data accessibility, GSA has introduced a new data retrieval system tailored to meet the diverse needs of researchers. This advanced retrieval interface enables users to conduct sophisticated queries across multiple search fields, encompassing organism, file type, sequencing platform, sequencing strategy, and numerous other criteria. The system further empowers users by allowing them to construct intricate queries using logical operators, facilitating the precise targeting and swift retrieval of desired data. Once search results are obtained, users are presented with a range of filtering options to refine their search outcomes even further. This flexibility ensures that researchers can quickly zero in on the most relevant data sets. Moreover, GSA offers a convenient batch download tool, enabling the seamless acquisition of search results in various formats, significantly streamlining the process of data collection and analysis.

In addition, GSA has introduced a sequence taxonomic analysis tool, derived from the Sequence Taxonomic Analysis Tool (STAT) [15]. This tool offers a comprehensive platform for the taxonomic classification analysis of the submitted data, complemented by intuitive web-based visualizations of the analysis results. By leveraging this service, researchers can gain valuable insights into the quality and validity of the data from a species-centric perspective. This, in turn, empowers them to make informed decisions regarding data selection and utilization, ultimately facilitating the efficient acquisition of the required data for their studies.

#### Genome Sequence Archive for Human

GSA-Human is a data repository dedicated to archiving, managing, and sharing raw sequence data from human biomedical research endeavours. GSA-Human provides two types of data access modalities: open access and controlled access. Open access allows unrestricted data downloads, whereas controlled access requires prior authorization from the Data Access Committee (DAC), as designated by the data submitter. Furthermore, GSA-Human has formulated comprehensive policies governing the sharing of human genetic resources data (https://ngdc.cncb.ac.cn/gsa-human/policy), providing users with clear guidelines on data submission procedures, management practices, and sharing protocols. These measures ensure both the security of sensitive data and its efficient utilization by the scientific community.

In recent years, the data volume in GSA-Human has experienced an exponential surge, with the current magnitude being a staggering sevenfold increase compared to 2021 levels. Notably, the most rapid expansion of 1 petabyte (PB) was achieved within a mere two days, posing significant challenges to data quality control procedures. To alleviate the pressure and improve data processing efficiency, we have upgraded the algorithm of the quality control process for FASTQ files by optimizing the data structure and minimizing memory consumption. In data quality control, for example, READ duplication detection typically uses a hash table to store all encountered READ names. However, as the file size increases, this approach leads to significant memory consumption, making it difficult for the quality control program to effectively process compressed files larger than 200 GB. The Bloom Filter is introduced in our new algorithm as an alternative to the hash table for marking READ duplication. It utilizes multiple bits to represent the presence of READ names, rather than storing their full information, thereby substantially reducing memory requirements and significantly enhancing the ability of the quality control program to handle large-scale datasets. The results indicate that the new algorithm reduces memory usage by 15 times, enabling the processing of terabyte-scale compressed files, and has been successfully integrated into GSA-Human and other relevant systems for processing raw sequence data.

In addition, GSA-Human has devised an intricate scoring system to evaluate the degree of sharing exhibited by the published datasets. Open access datasets automatically receive the full scores, while for controlled access datasets, a comprehensive evaluation incorporates various factors related to the DAC’s (Data Access Committee) handling of data access requests. These factors include the percentages of processed and approved requests, and the average processing time for the requests, which together contribute to determining the final score. This scoring methodology offers an intuitive measure of dataset sharing levels, furnishing data requesters with clear and understandable insights. It is our hope that such a scoring system will foster a culture of data sharing and encourage the reuse of scientific data. The scoring details for the public datasets are available online (https://ngdc.cncb.ac.cn/gsa-human/browse/).

#### Open Archive for Miscellaneous Data

OMIX, built adhering to the FAIR principles[18], stands as a versatile data repository, dedicated to gathering, publishing, and sharing scientific data for biological researchers. Currently, OMIX supports ten primary data types, including 32 sub-categories that span functional genomics, proteomics, metabolomics, and more (Table S1), underscoring its comprehensive capabilities and adaptability.

OMIX provides two distinct data access modalities: open access and controlled access, both available for human genetic resource data, while data from other species is limited to open access. In managing and authorizing controlled data access, OMIX employs a mechanism similar to GSA-Human but simplifies the “request and approval” process. This allows data requesters to easily submit applications and provides data providers with a one-click approval feature, thus reducing processing time and enhancing the efficiency of data acquisition.

As a significant member of the GSA family, OMIX expands the scope of data intake and enhances data diversity. By linking to other member databases via BioProject IDs, it creates multi-omics datasets centered around raw sequencing data from GSA or GSA-Human, providing robust support for advancing scientific research. For instance, BioProject PRJCA006118 contains 25 paired datasets primarily focused on transcriptomics and single-cell sequencing, while also encompassing proteomics, clinical data, and imaging data. With a total data volume reaching 9.21 TB, this resource has supported six published research articles.

#### Open Biomedical Imaging Archive

OBIA, the new member of the GSA family, serves as an innovative data repository meticulously crafted to store and share multimodal medical imaging alongside its clinical data. It boasts a dual-pronged approach to data accessibility, encompassing both open and controlled access mechanisms, catering to varying user needs and security requirements. This feature underscores OBIA’s pivotal role in fostering collaborative research by bridging the gap between diverse data modalities and facilitating the synthesis of insights for advanced medical discoveries.

The primary challenge in constructing a medical image database is ensuring robust privacy protection. Images may contain protected health information (PHI) and require appropriate processing before archived to minimize the risk of patient privacy breaches. OBIA addresses this by providing a unified de-identification and quality control mechanism based on the Digital Imaging and Communications in Medicine (DICOM) standard. The key elements and rules we adopted include: 1) clean pixel data, 2) clean descriptors, 3) retain longitudinal temporal information modified dates, 4) retain patient characteristics, 5) retain device identity, and 6) retain safe private tags. OBIA utilizes the Radiological Society of North America (RSNA) MIRC clinical trial processor (CTP) (https://mircwiki.rsna.org/index.php?title=MIRC_CTP) to remove or blank certain standard tags containing or potentially containing PHI. For private tags, we employ PyDicom (https://pypi.org/project/pydicom/) to retain attributes that are purely numeric. Furthermore, images with PHI in their pixel data are rigorously filtered to safeguard patient privacy.

In addition to providing web interfaces for data query and browsing, OBIA offers innovative image retrieval capabilities[19]. The retrieval model is based solely on image features and uses the Hamming distance to return visually similar images. Users can upload images to search the database for the top 30 most similar images, significantly enhancing the user experience.

### Data submission

The GSA family is equipped with the Single sign-on (SSO) service, a user access control system that enables seamless authentication across multiple databases with a single, unified ID and password. To submit data to the GSA family, users are required to initially register an account within the SSO, and log into the specific database where they intend to initiate their submissions.

Each database within the GSA family provides a convenient web-based submission service that defines a series of metadata templates tailored to the characteristics of the accepted data, facilitating the collection of the required information. Take GSA-Human as an example, it offers specialized templates for human-related researches, encompassing diverse domains such as disease, cohort, cell line, clinical pathogen and human-associated metagenome studies. The templates cover essential attributes required for the data objects, spanning sample details, experimental conditions, and sequencing strategies, while also allowing users to incorporate custom attributes, greatly enhancing adaptability to personalized research requirements.

Moreover, each database within the GSA family has quality control and expert curation processes to ensure the accuracy and reliability of the submitted data. These processes focus on verifying the integrity, validity, and consistency of the metadata and data files, as well as identifying and removing personal privacy information, thereby ensuring the preservation of high-quality data for future reference and analysis.

### Data mirroring

The INSDC maintains the largest collection of raw sequence data that are not readily retrievable for many researchers in China, mainly due to the large volume and limited network bandwidth. The three member databases of INSDC, namely SRA, DDBJ, and ENA, exchange and share data on a daily basis. As of February 2024, there is a total of nearly 28 PB of published data in SRA, while the data volume in GSA is 5.3 PB. In an effort to facilitate the seamless sharing and reuse of this global data, GSA has developed a system that mirrors the raw sequence data of INSDC sourced from NCBI. Metadata including BioProjects, BioSamples and SRAs are synchronously downloaded, parsed and stored locally, and the sequence files are also mirrored in accordance with the metadata. The full metadata and the newly released sequence files since April 2022 are mirrored and synchronized daily, ensuring the data consistency with INSDC. Furthermore, all the mirrored data are integrated into GSA and powered with the retrieval, browse, and download functions, offering users with convenient ways to access the data. The integrated GSA, which combines data from multiple sources, offers a broader range of information compared to other omics datasets, thereby providing users with a more comprehensive and enriched database.

### Data statistics

As of December 2024, the GSA family has archived 28,838 datasets, totalling over 59.98 petabytes (PB) of data. From 2015 to 2024, there was a remarkable growth of data in the family. The data volume has experienced exponential growth, and the diversity of data types has notably expanded (**Figure 2, Table 1**). Its data accessions have been cited in 4,351 research articles across 729 scientific journals (https://ngdc.cncb.ac.cn/gsa/statistics?active=journals). For detailed statistics on data submissions by each database.

**Table 1.** Data submissions within the GSA Family.

| Data Submitted Item | GSA | GSA-Human | OMIX | OBIA | Total |
| --- | --- | --- | --- | --- | --- |
| No. of datasets | 16,001 | 6602 | 6229 | 6 | 28,838 |
| No. of open-access datasets | 16,001 | 1559 | 2925 | 1 | 20,486 |
| No. of controlled-access datasets | / | 5043 | 3304 | 5 | 8352 |
| No. of BioProjects | 13,583 | 5888 | 4035 | 3 | 23,509 |
| No. of Individuals | / | 582,817 | / | 1094 | 583,911 |
| No. of Samples | 889,841 | 847,958 | / | / | 1,737,799 |
| No. of Experiments | 891,329 | 952,690 | / | / | 1,844,019 |
| No. of Runs | 965,023 | 1,248,968 | / | / | 2,213,991 |
| No. of Files | 1,744,888 | 1,943,316 | 32,094 | 2,012,007 | 5,732,305 |
| Total submitted file size (TB) | 19,722.24 | 41,605.12 | 90.97 | 0.39 | 61,418.72 |
| No. of support publications | 2497 | 1037 | 974 | / | 4351 |

*Note:* All statistics regarding data submissions were derived from the GSA family as of December 2024. “/” means not applicable. TB, terabyte. Due to articles referencing serial accessions from multiple databases simultaneously, the total article count is lower than the sum of individual article counts in each database.

**Figure 2.**
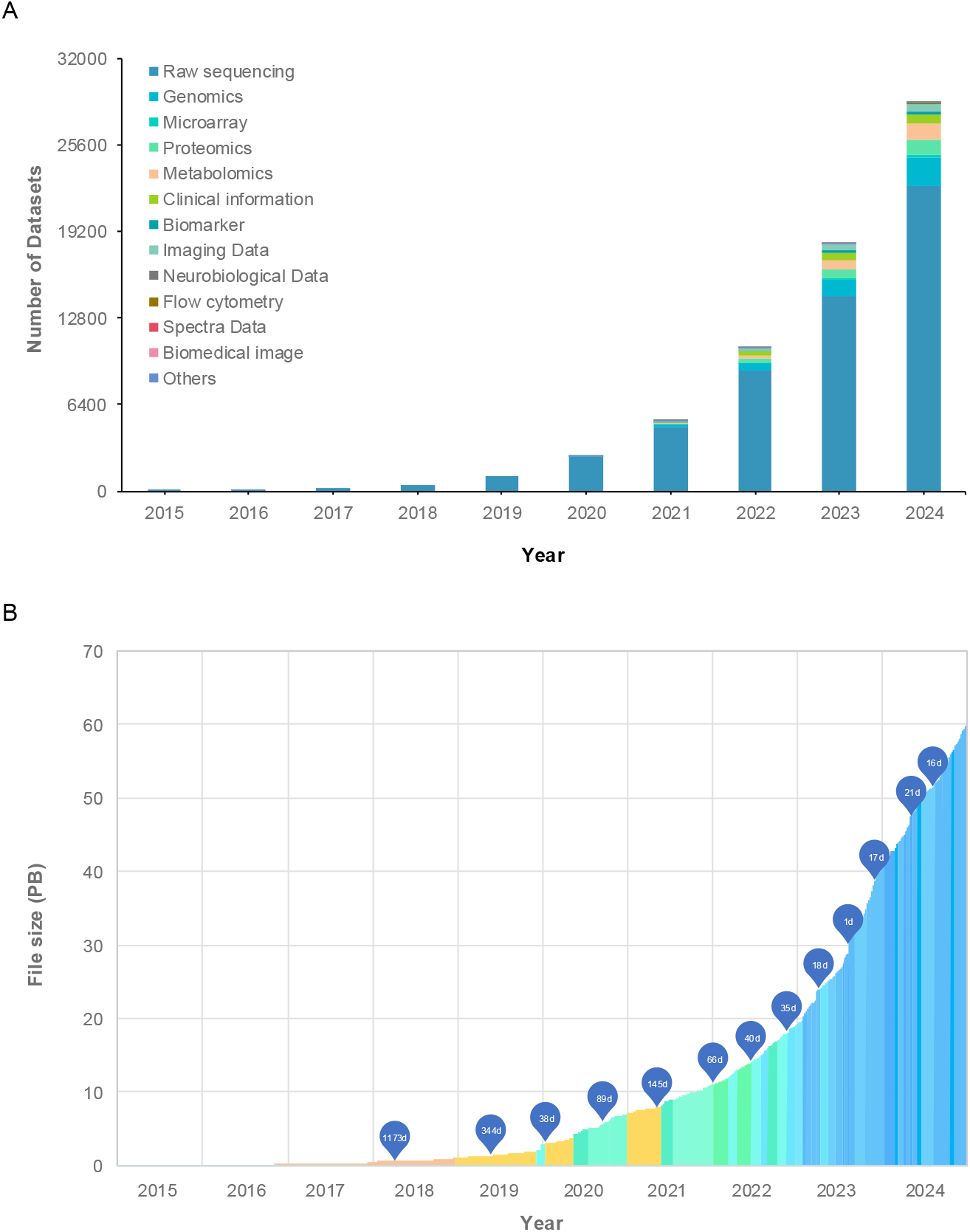
Data statistics of the GSA family. **A**. Number of datasets accumulated from 2015 to 2024 from GSA family, with thirteen major data types indicated. **B**. Increase in the volume of submitted raw sequence data over time, with the time required to accumulate each PB of data indicated. All statistics were derived from GSA and GSA-Human as of December 2024. The label inside the bubble indicates the number of days required for the data volume to increase by 1 PB. PB, petabyte; d, days.

In terms of data sharing, the GSA family offers two main access modalities: open and controlled access. Through open access, the database provides researchers with unrestricted access to datasets. As of December 2024, the GSA family has published 14,557 open access datasets, amounting to 7.45 PB of data shared via FTP, HTTP, with a total of 166,549,212 downloads. For controlled data sharing, the GSA family has published 4,756 controlled access datasets, receiving a total of 9,383 access requests from 4,829 requesters, with 3,242 requests approved, resulting in nearly 10 PB of data downloads (For detailed statistics on data sharing for each database, see Table 2). Regarding international data mirroring, as of December 2024, GSA has mirrored 783,566 projects, 41,867,651 samples, 32,507,936 experiments, and 34,456,823 runs, as well as over 10.8 PB of sequence files from NCBI.

**Table 2.**
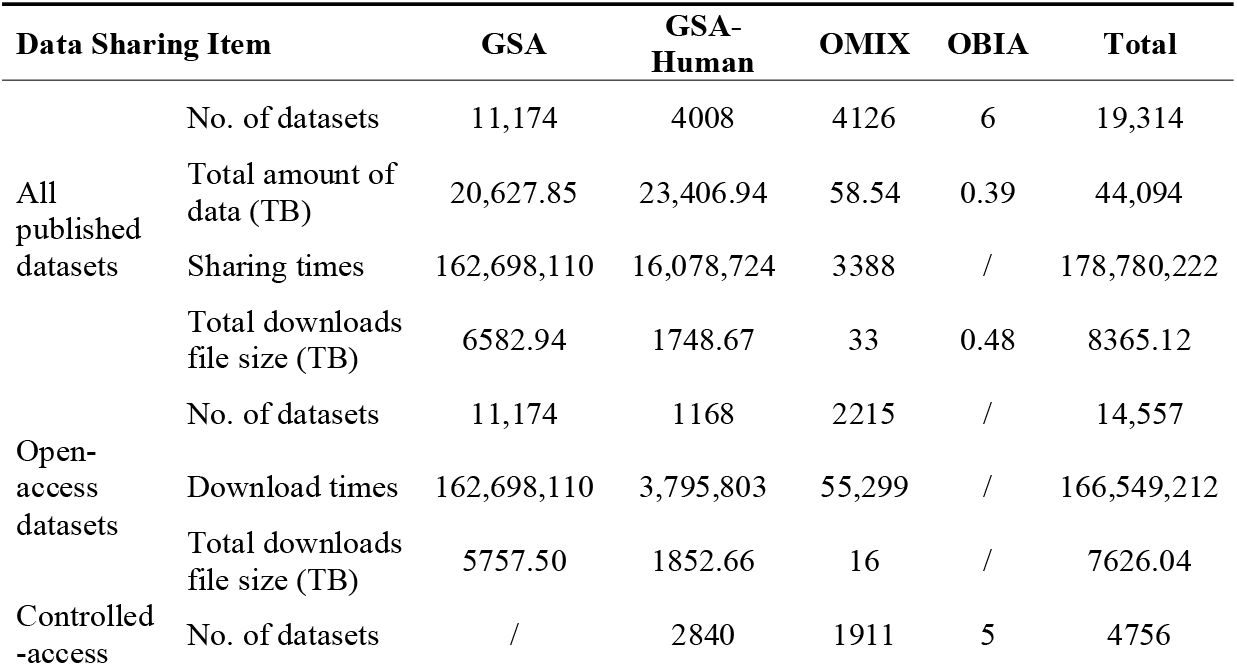
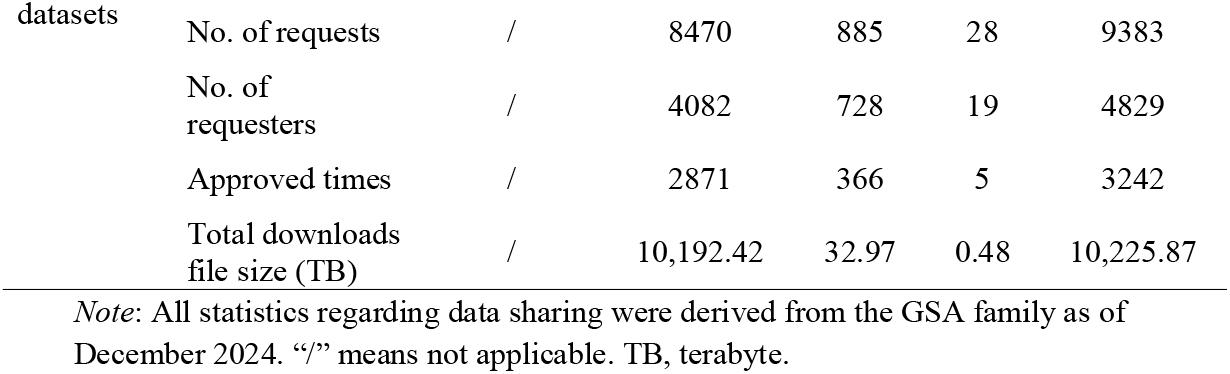
Data sharing within the GSA family.

### Future direction

The GSA family has continuously enhanced its archival databases for raw sequence, multi-omics, and multimodal data, streamlining data submission, management, and reuse. Unlike conventional archives, this four-member family supports a broader spectrum of data types across diverse species and research fields. Its innovative management approach, cantered on raw omics data, enables efficient multi-omics pairings and delivers a comprehensive suite of services to users.

Future efforts will be concentrated on continuous optimizations of data models and management processes to meet evolving user demands and rapidly increasing data volumes. An advanced cloud-based data storage architecture, as long as a suit of innovative tools, will be established to improve the efficiency of storing and mining the omics big data. To tackle the challenges presented by the exponential surge in data, a more automated data management system will be established to streamline and automate the processes from data submission, reception, quality control, archive, publication to update, so as to improve data processing efficiency.

For the secure and sustainable management of human genetic resource data, a comprehensive strategy will be employed. This includes the implementation or enhancement of data classification frameworks, data management mechanisms, reliable backup and recovery systems, as well as advanced anonymization and encryption technologies. In parallel, the exploration of a secure cloud-based data analysis model is anticipated, aiming to mitigate privacy risks associated with the direct download of raw data. By utilizing a controlled storage environment for sensitive files, this model significantly reduces the potential for data exposure. Furthermore, personal identification techniques based on genomic sequences will also be considered.

Additionally, active efforts are underway to promote global collaboration aimed at advancing data standards, technologies, and methodologies. Concurrently, work is being conducted to achieve full mirroring and synchronization of data from the International Nucleotide Sequence Database Collaboration (INSDC), along with the integration of additional global omics database resources. These initiatives are designed to support the widespread sharing and utilization of global biodiversity and health-related big data.

## Data availability

GSA is open accessible at https://ngdc.cncb.ac.cn/gsa/.

GSA-Human is freely accessible at https://ngdc.cncb.ac.cn/gsa-human/.

OMIX is freely accessible at https://ngdc.cncb.ac.cn/omix/.

OBIA is freely accessible at https://ngdc.cncb.ac.cn/obia.

## CRediT author statement

**Sisi Zhang**: Investigation, Methodology, Data curation, Writing - original draft, Writing – review & editing. **Xu Chen**: Software, Writing - original draft. **Enhui Jin**: Software, Investigation, Data curation, Writing - original draft. **Anke Wang**: Software, Writing - original draft. **Tingting Chen**: Investigation, Methodology, Data curation, Writing - original draft. **Xiaolong Zhang**: Software, Writing - original draft. **Junwei Zhu**: Software. **Lili Dong**: Data curation. **Yanling Sun**: Data curation. **Caixia Yu**: Data curation. **Yubo Zhou**: Software. **Zhuojing Fan**: Software. **Huanxin Chen**: Resources. **Shuang Zhai**: Resources. **Yubin Sun**: Resources. **Qiancheng Chen**: Resources. **Jingfa Xiao**: Conceptualization. **Shuhui Song**: Conceptualization. **Zhang Zhang**: Conceptualization. **Yiming Bao**: Conceptualization. **Yanqing Wang**: Conceptualization, Investigation, Methodology, Software, Writing - review & editing. **Wenming Zhao**: Conceptualization, Methodology, Writing - review & editing, Supervision. All authors have read and approved the final manuscript.

## Acknowledgments

This work was supported by National Key R&D Program of China (Grant Nos. 2023YFC2605702 to WZ, 2023YFC2604402 to YW, 2021YFF0703701 to YB); Strategic Priority Research Program of the Chinese Academy of Sciences (Grant No. XDA0460203); National Natural Science Foundation of China (Grant Nos. 32100511 to YW, 32170678 to WZ; 32030021 to ZZ; 92374201 to SS); Strategic Priority Research Program of Chinese Academy of Sciences (Grant No. XDB38050300 to WZ).

