## Supplemental Table 1 for "The GSA Family in 2025: A Broadened Sharing Platform for Multi-Omics and Multimodal Data"

### Supplementary Materials

#### Table S1 Data type of OMIX database

| **Data Type** | **Sub Type** | **Datasets** | **Files** | **File Size (TB)** |
| --- | --- | --- | --- | --- |
| Genomics | Methylation Profiling by NGS | 2,073 | 10177 | 20.35 |
|  | Expression Profiling by NGS |  |  |  |
|  | Non-coding RNA Profiling by NGS |  |  |  |
|  | Genome Binding/Occupancy Profiling by NGS |  |  |  |
|  | Chromatin Accessibility Profiling by NGS |  |  |  |
|  | Other Type of Genomic Data |  |  |  |
| Microarray | Methylation profiling by Array | 282 | 1,425 | 0.26 |
|  | Expression Profiling by Array |  |  |  |
|  | Non-coding RNA Profiling by Array |  |  |  |
|  | Genome binding/occupancy profiling by Array |  |  |  |
|  | Other type of Microarray Data |  |  |  |
| Proteomics | Protein 3D Structure Data | 1048 | 7861 | 39.19 |
|  | Proteomic Data by Mass Spectrometry |  |  |  |
|  | Other Type of Proteomic Data |  |  |  |
| Metabolomics | Metabolome Data by Mass Spectrometry | 1231 | 9776 | 19.81 |
|  | Lipidome Data by Mass Spectrometry |  |  |  |
|  | Other Type of Metabolome Data |  |  |  |
| Clinical Information | Demographic Data | 638 | 874 | 0.92 |
|  | Clinical Research Data |  |  |  |
|  | Other Type of Clinical Information |  |  |  |
| Biomarker | Genetic Biomarkers | 535 | 675 | 0.90 |
|  | Karyotype Biomarkers |  |  |  |
|  | Protein Biomarkers |  |  |  |
|  | Chemical Biomarkers |  |  |  |
|  | Condition-specific Biomarkers |  |  |  |
| Imaging | Magnetic Resonance Imaging | 228 | 772 | 4.65 |
|  | Behavioral Video Data |  |  |  |
|  | Other Type of Image Data |  |  |  |
| Neurobiological Data | / | 30 | 60 | 0.75 |
| Flow cytometry | / | 51 | 180 | 0.13 |
| Spectra Data | Raman Spectra Data | 10 | 15 | 0.02 |
|  | Other type of Spectra Data |  |  |  |
| Others | / | 103 | 279 | 3.99 |

*Note*: All statistics regarding data sharing were derived from the OMIX as of December 2024. “/” means not applicable. TB, terabyte. Prior to 2022, OMIX allowed the submission of multiple data types within a single dataset. To enhance data usability and accessibility, OMIX subsequently refined its data categories, expanding them to 10 major categories and 32 subcategories. Datasets previously marked as "Other" by users during submission include imaging data, molecular spectra, protein sequences, CT scans and biomarker data. Following system enhancement, these data types have been incorporated into the expanded OMIX categories, and the "Other" classification no longer appears in subsequent submissions.
